# Cinemeducation improves early clinical exposure to Inborn Errors of Metabolism

**DOI:** 10.1101/2022.08.30.505958

**Authors:** Atanu Kumar Dutta, Aroma Oberoi, Kalyan Goswami, Jyoti Nath Modi, Parmod Kumar Goyal, Sangeetha Samuel, Tanushree Mondal, Sibasish Sahoo, Amit Pal

## Abstract

**Background:** Cinemeducation has been shown to be an effective tool to help the students develop humanistic skills. However, there is a dearth of studies to find out if this can also be utilized to improve the interest and satisfaction of students learning about rare diseases such as the Inborn Errors of Metabolism. The aim was to introduce cinemeducation as part of early clinical exposure in teaching the inborn errors of metabolism for first-year MBBS students in the Department of Biochemistry. The Objectives were 1) to develop, validate and assess the feasibility of structured modules on Cinemeducation in teaching the Inborn Errors of Metabolism, 2) to study the impact of Cinemeducation on the satisfaction, interest, and academic performance of first-year MBBS students and 3) to study the perception of students and faculty towards Cinemeducation

**Methods:** This was a quasi-experimental, cross-over study involving 100 MBBS students in the Department of Biochemistry, AIIMS Kalyani. Movie clips prepared from Lorenzo’s Oil (for Lipid metabolism) and Extraordinary Measures (for Lysosomal Storage Disease) and corresponding paper-based cases; questionnaires and feedback forms were validated by expert group review. Analysis of quantitative data was performed in GraphPad, MS Excel and MAXQDA2022 software.

**Findings, Discussions and Conclusions:** Students’ academic performance was found to be improved in the groups where the Extraordinary Measures movie was used for Cinemeducation. Both learners and facilitators were satisfied with Cinemeducation. Cinemeducation was effectively introduced for teaching Inborn Errors of Metabolism in the Department of Biochemistry.

## Background

Inborn errors of metabolism (IEM) are amenable to good outcomes with early diagnosis and management. However, because of the non-availability of clinical material for these rare diseases, undergraduate medical students in India typically do not get exposed to them. The absence of patient exposure is an obstacle for students developing an interest in learning these topics and also in understanding their psychosocial aspects. Therefore, the facilitators also face difficulty to include these topics in the formative and summative assessments thereby further diminishing the interest of the learners. Thus, the Indian Medical Graduates are lacking in the understanding of their role in the early diagnosis of inborn errors of metabolism through clinical suspicion and appropriate referral and also diagnostic / management challenges unique to these diseases. This leads to a delay in their diagnosis and adverse outcomes.

Though Medical Genetics is currently transforming the practice of Medicine; in India, it is patchily represented in the undergraduate (UG) medical curriculum. Therefore, there is an urgent need to include more elements of Medical Genetics in the UG teaching-learning activities.^1^ Medical Biochemistry includes the theoretical framework of genetics, diagnostic techniques, and mechanism of genetic diseases. I intended to use an unconventional teaching-learning methodology namely movie clips to facilitate students learning in Genetics. Using movies in medical education has been shown to be an effective tool, especially for training students in professionalism, ethics, and critical thinking.^2^ This approach is already in use abroad in the form of Film Club (Mediaomics).^3,4,5^ The use of movie clips has already been shown to be effective in medical education in a number of studies involving teaching in Neuropsychiatry and Infectious diseases.^6,7,8,9^ Ideally, there is a need for a blended approach utilizing the strengths of movie clips and conventional teaching.^10^ Moreover, the effectiveness of using audio-visual aids in building a conceptual framework for medical students has been well documented.^11,12^ Arguably the best movie where Biochemistry itself is the central character is “Lorenzo’s Oil”.^13,14^ This movie can be an excellent tool for teaching Inborn Errors of Metabolism.

### Aim

To introduce cinemeducation as part of early clinical exposure in teaching the Inborn Errors of Metabolism for first-year undergraduates in the Department of Biochemistry.

### Objectives

i. To develop, validate and assess the feasibility of structured modules on Cinemeducation in teaching the Inborn Errors of Metabolism.
ii. To study the perception of students towards Cinemeducation in teaching the Inborn Errors of Metabolism for the first-year MBBS students.
iii. To study the perception of faculty towards Cinemeducation in teaching the Inborn Errors of Metabolism for the first-year MBBS students.
iv. To study the impact of Cinemeducation on the satisfaction, interest, and academic performance of first-year MBBS students.

## Methods

### Study design

Quasi-experimental, cross-over, non-randomized, interventional study.

### Setting

Department of Biochemistry, AIIMS Kalyani

### Participants

Year 1 MBBS students from the 2020 batch.

### Sample size

100, divided into four equal groups of 25 students each.

### Sampling/ Grouping strategy

Complete enumeration. The whole batch of students was taken as a sample. No randomization was done. Students are assigned to four groups (A, B, C, D) of 25 students each, according to their class roll numbers and attendance in the sessions.

Movie clips were prepared from Lorenzo’s Oil (for Adrenoleukodystrophy) and Extraordinary Measures (for Pompe Disease) and corresponding paper-based cases; Pre and Post-test questionnaire, Student’s feedback form including Likert scale and open-ended questions, Faculty focused group discussion.

### Validation of tools

Movie clips were selected and then finalized by a Departmental committee; questionnaires were validated by an expert group.

### Study implementation

Lorenzo’s Oil and Extraordinary Measures movies were purchased from the United States of America. Showing movies on the educational campus to selected students is exempted from Public Performance Rights under the face-to-face teaching exemption at 17 U.S.C. §110(1) (https://guides.uflib.ufl.edu/copyright/video). Lorenzo’s Oil movie has also been given access to be used for educational and research purposes by the US Library of Congress (https://www.loc.gov/item/jots.200161376/).

Description of Intervention: Students belonging to groups A and B were shown clips from Lorenzo’s Oil and a paper-based case derived from Extraordinary Measures. Students belonging to groups C and D were shown clips from Extraordinary Measures and a paper-based case derived from Lorenzo’s Oil (figure 1). Movie clips were selected through a Departmental meeting based on their relevance to the topic, limited time available in the class, and cultural acceptability (supplementary table 1).

**Figure 1:**
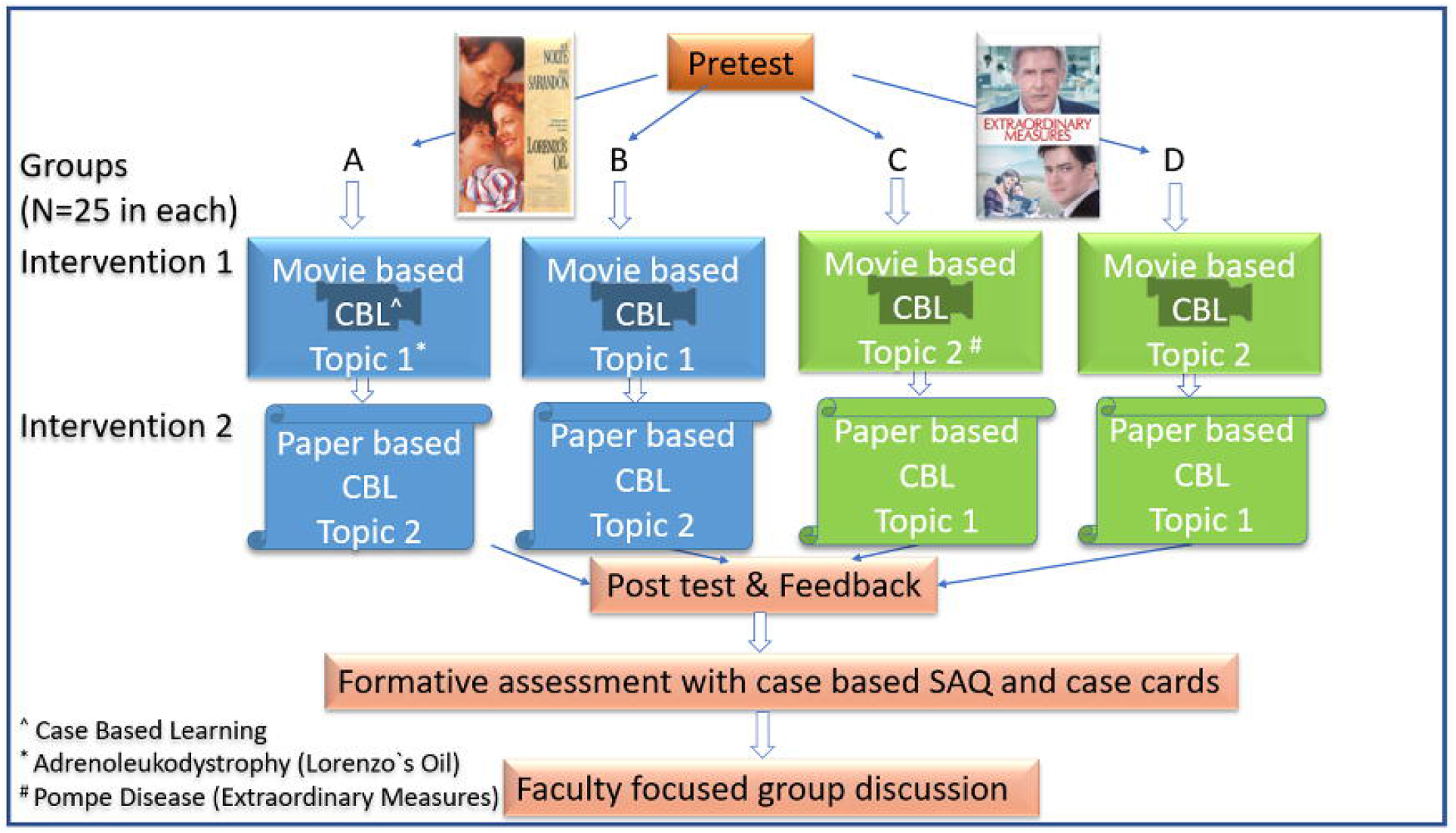
Intervention Flow Diagram

### Data collection process

Pre-test, Post-test, and students’ feedback were collected in paper and then transcribed in Excel sheet. A focused Group Discussion among the faculty was recorded and the audio was transcribed by the Maestra suite web interface (https://maestrasuite.com/). One case-based question on Lysosomal storage disease was included in the formative assessment.

## Data analysis

### Quantitative

Analysis of quantitative data was performed in GraphPad Prism 9 software. Consensus calculation for the Likert scale data was performed in MS Excel according to Tasle and Wierman.^15^ The first distribution of the data was checked for normality. As the data did not follow normality nonparametric methods were chosen for statistical analysis. To test for a difference in the mean (or median) Wilcoxon matched-pairs signed ranked test was used for paired data and the Mann Whitney test was used for unpaired data.

### Qualitative

Thematic analysis of the student’s feedback and Faculty FGD was performed using MAXQDA2022 software. The identified themes were used to perform a force field analysis of the utility of Cinemeducation to supplement Case Based Learning in Biochemistry.

### Ethics

Written informed consent was taken from the students following a brief discussion. As we have employed a crossover design hence all the groups of students had exposure to both the movie as well as the conventional CBL in one topic each.

## Findings

We were able to conduct the Cinemeducation & CBL sessions as planned. Apply the pre and post-tests and collect feedback from the learners following the validation of the tools and modules by peer and expert feedback. We also included one case-based question on Pompe disease in the formative assessment. A focused group discussion was conducted for the facilitators.

The descriptive statistics of the pre-test post-test and formative assessment data of the different intervention groups are presented in supplementary table 2. The distribution of these data does not follow a normal distribution as demonstrated in the Q-Q plot in supplementary figure 1. The Violin plot of the pre-test and post-test scores of the different interventional groups is shown in figure 2. The post-test score was significantly improved from the post-test score in all the groups. However, when the post-test score of topics taught using Cinemeducation or CBL was compared the difference was only significant when the Extraordinary Measures movie was used for the Pompe Disease topic. There was no significant difference between groups (A+B) and (C+D) in the formative assessment before this intervention (figure 3a). However, group (C+D) performed significantly well in the formative assessment question compared to group (A+B) (figure 3b). Interestingly this remained valid for questions belonging to other topics as well (figure 3c). The perception of the learners about cinemeducation based on their Likert scale feedback is presented in table 1. The results of the thematic analysis of learners and focused group discussion of the facilitators are presented in a triangulation manner in supplementary table 3 and summarized in figure 4 through force field analysis. Students feedback is quoted in non-italicized and facilitator’s feedback is quoted in italicized letters in supplementary table 3.

**Figure 2:**
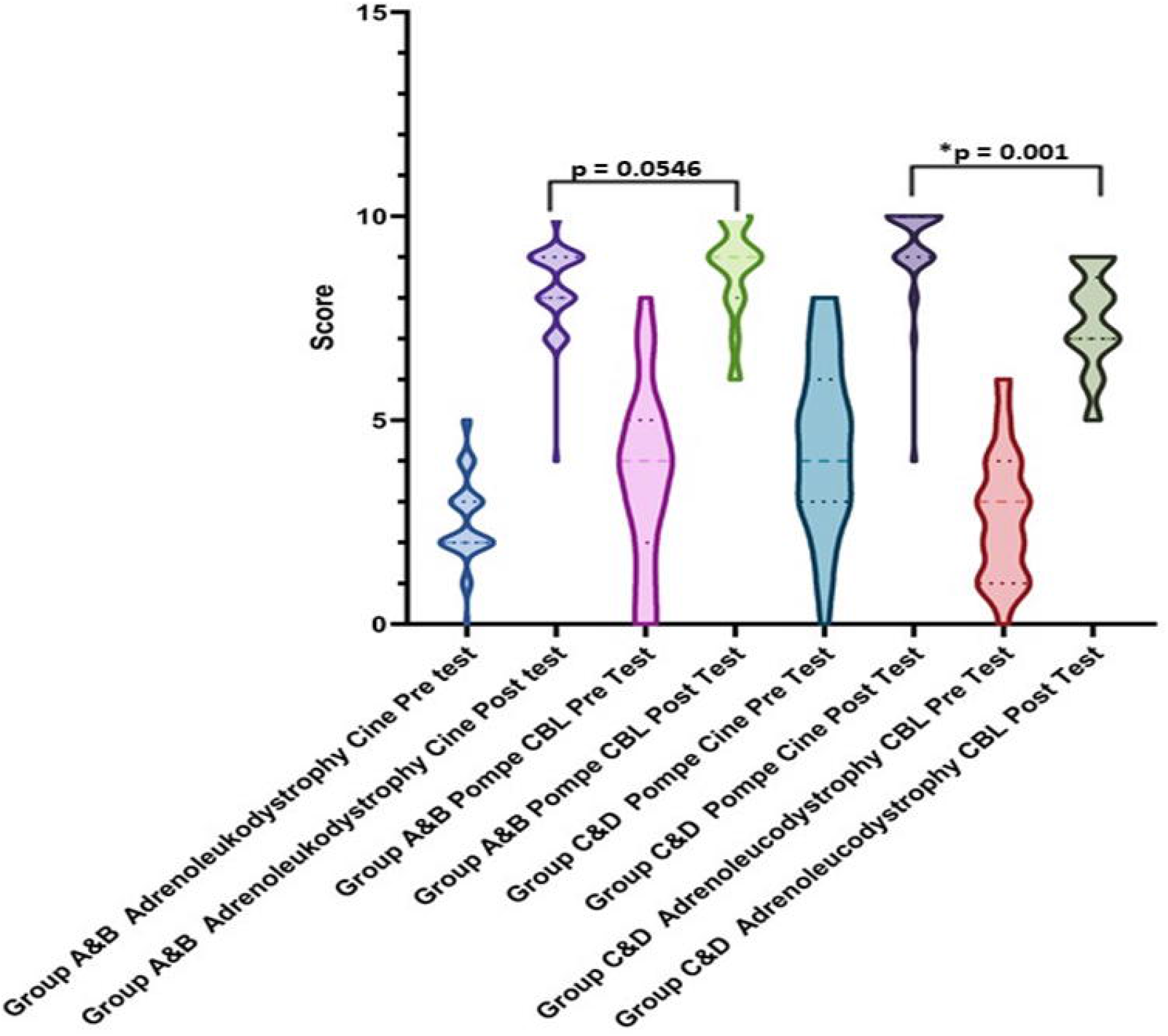
Violin plot of pre and post-test scores of different intervention groups. Post-test scores within groups were analyzed by the Wilcoxon matched-pairs signed ranked test. Two-tailed p-value was considered to be significant if <0.05.

**Figure 3:**
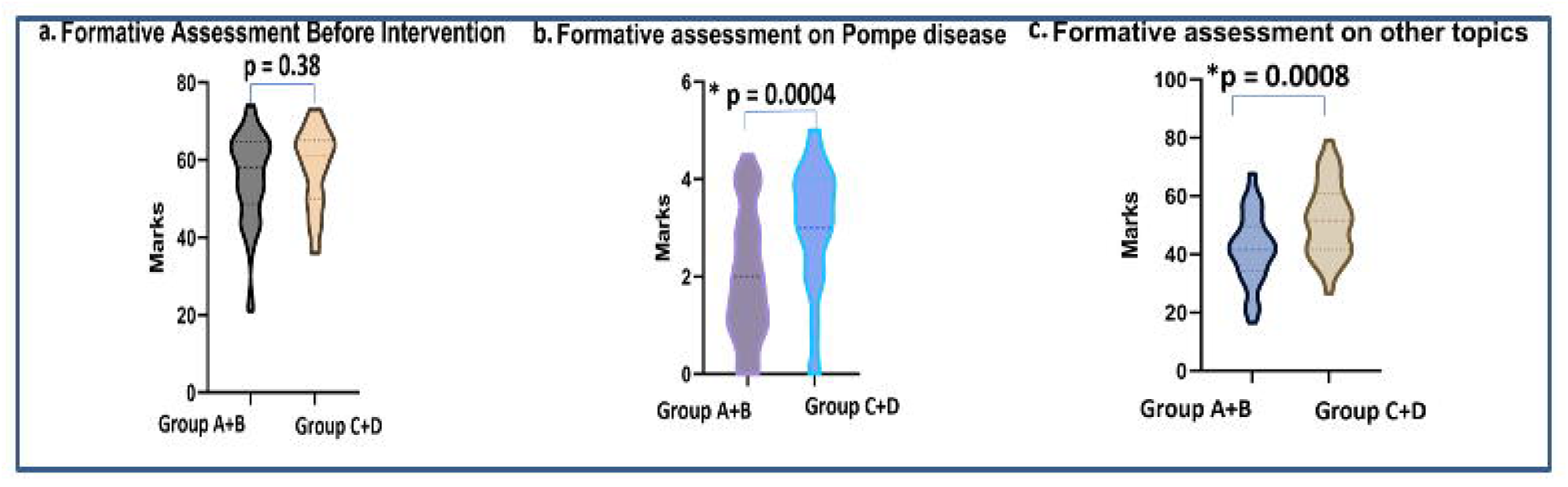
Violin plot of formative assessment scores of the different groups before (a) and after (b & c) intervention. Formative assessment scores between groups were analyzed by Mann Whitney test. Two-tailed p-value was considered to be significant if <0.05.

**Table 1.** Perception of learners about cinemeducation

| Sl. No. | Statement | Strongly agree and agree (%) | Consensus |
| --- | --- | --- | --- |
| 1 | Use of movie clips increased my interest in Inborn Errors of Metabolism when compared to conventional case-based learning (CBL) | 100 | 0.86 |
| 2 | I am satisfied with the use of movie clips for learning inborn errors of metabolism as compared to conventional CBL | 98 | 0.80 |
| 3 | Movie clips can be a good option to supplement the teaching-learning process | 98 | 0.81 |
| 4 | Movie clips can be a good option to complement the teaching-learning process | 81 | 0.62 |
| 5 | Choice of movies for the purpose was relevant | 99 | 0.84 |

**Figure 4:**
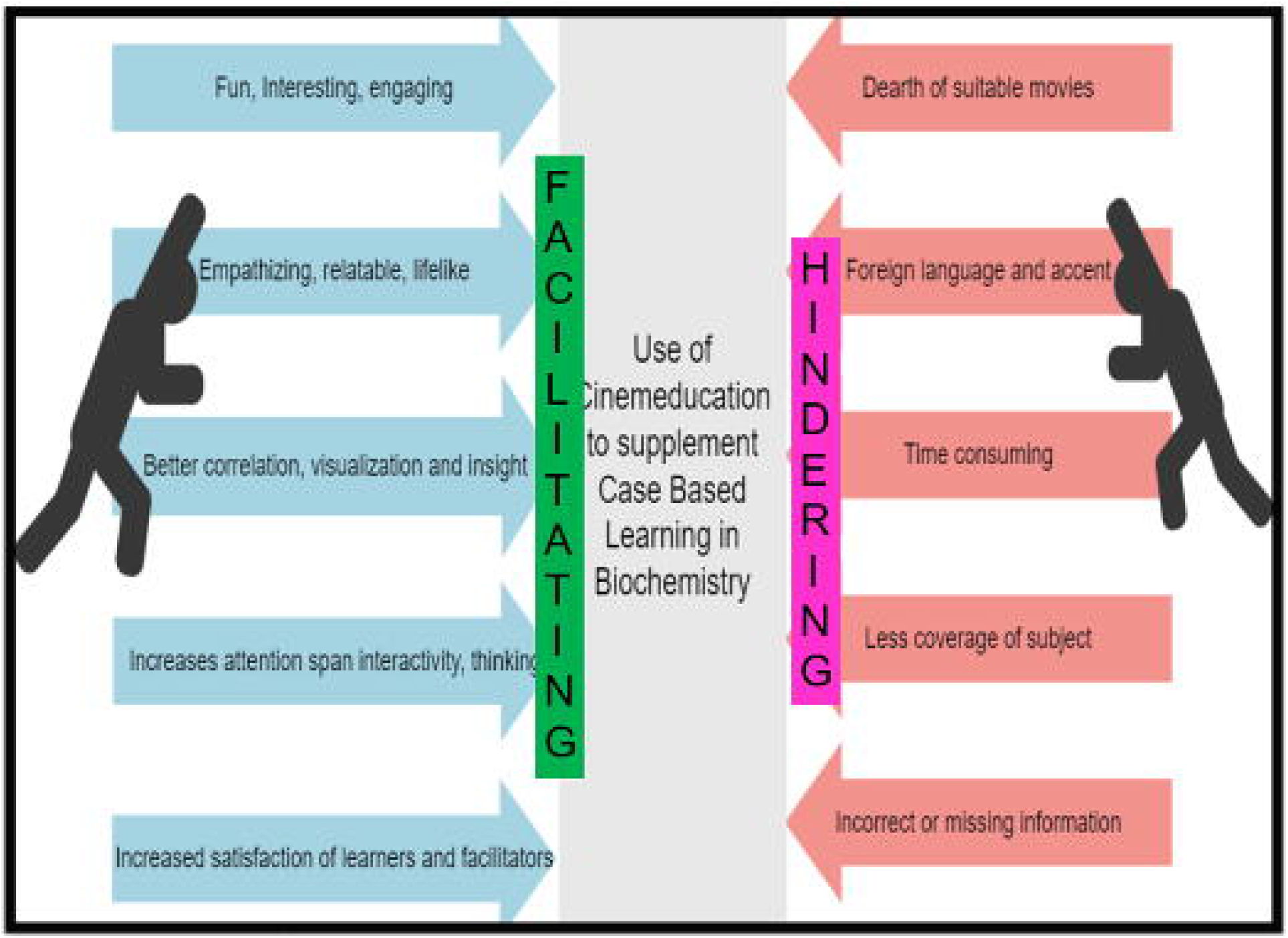
Force Field Analysis

## Discussion

The results showed that the success of Cinemeducation to be more effective than conventional CBL is critically dependent on the movie used. Here though Lorenzo’s Oil was heavier in terms of subject content; students liked Extraordinary Measures more partly due to the star cast in this movie. Extraordinary Measures movie was also more effective in teaching Pompe Disease as evident by better post-test scores compared to conventional CBL and better performance in the formative assessment.

In general, the learners strongly agreed that Cinemeducation increased their interest and satisfaction with learning Inborn Errors of Metabolism when compared to traditional CBL and that the choice of movies was relevant. However, they felt movie clips should be used to supplement the teaching-learning process and not complement the same. Taken together the facilitating factors much outweigh the hindering factors for the application of Cinemeducation.

These results clearly demonstrated that Cinemeducation strongly aroused interest in the learners and the satisfaction of both the learners and facilitators. Compared to the previous studies our learners found some difficulty in following clips from Lorenzo’s Oil.^13,14^ Possible reasons for this could be the language barrier/ the Italian accent present in this movie and relatively less well-known actors in this movie. Our study supports earlier findings that Cinemeducation helps to improve students’ motivation to learn and improve learning of the Humanistic aspects of medicine.^16,17^ Limitations of this study include the lack of long-term follow-up results and the inclusion of only the Pompe disease in the formative assessment. Adrenoleukodystrophy could not be included in the formative assessment as it was not in the must-know area. The study demonstrated that Cinemeducation is an effective unconventional strategy to increase the learner’s interest and satisfaction in learning Inborn Errors of Metabolism. In the future, we would collect longitudinal follow-up data from the learners and facilitators to study its long-term effectiveness such as in developing critical thinking.^18^ We would also make attempts to produce short movies that can be used for Cinemeducation.

## Conclusions

In conclusion we were able to develop and validate two structured modules on cinemeducation for teaching the inborn errors of metabolism. We found it is feasible to use these modules for first-year MBBS students. This led to increased interest and satisfaction of the learners and their academic performance also improved in Pompe disease when Cinemeducation was used as the teaching-learning method. The faculty was also satisfied with these modules and they suggested making more such small movies to be used for Cinemeducation in Biochemistry in the future.

