## Supplementary data for "Cinemeducation improves early clinical exposure to Inborn Errors of Metabolism"

**Supplementary table 1**

**Movie Clips Used**

| Sl. No. | Topic | Movie | Clip timing | Description |
| --- | --- | --- | --- | --- |
| 1 | Adrenoleukodystrophy  (Full length 2:15:10) | Lorenzo`s Oil  (Total clip duration 39 minutes) | i) 03:37 – 06:27 | Disease prodrome |
|  |  |  | ii) 08:24 – 13:53 | Diagnostic workup |
|  |  |  | iii) 14:25 – 22:49 | Dietary and Immunological clinical trials |
|  |  |  | iv) 23:45 – 34:53 | Fatty acid manipulation with C18 MUFA Oleic Acid supplementation |
|  |  |  | v) 41:40 – 53:15 | Competitive inhibition of fatty acid elongase by C22 MUFA Erucic Acid supplementation |
| 2 | Pompe Disease  (Full length 1:46:00) | Extraordinary Measures  (Total clip duration 29 minutes) | i) 1:16 -12:16 | Clinical features and inheritance |
|  |  |  | ii) 8:30 - 13:55 | Acute exacerbation |
|  |  |  | iii) 15:50 – 17:36 | Theory of Enzyme Replacement Therapy |
|  |  |  | iv) 1:27:10 -1:33:20 | Sibling trial |
|  |  |  | v) 1:35:25 – 1:40:35 | First ERT trial |

Supplementary table 2

Descriptive statistics

| Sl. No. | Groups | Mean | Std. Deviation | Std. Error of Mean |
| --- | --- | --- | --- | --- |
| 1 | Group A&B Adrenoleukodystrophy Cine Pre-test (out of 10) | 2.6 | 1.0 | 0.1 |
| 2 | Group A&B Adrenoleukodystrophy Cine Post-test (out of 10) | 8.3 | 1.0 | 0.1 |
| 3 | Group A&B Pompe CBL Pre Test (out of 10) | 3.7 | 2.1 | 0.3 |
| 4 | Group A&B Pompe CBL Post Test (out of 10) | 8.7 | 1.2 | 0.2 |
| 5 | Group C&D Pompe Cine Pre-Test (out of 10) | 4.4 | 2.1 | 0.3 |
| 6 | Group C&D Pompe Cine Post Test (out of 10) | 9.3 | 1.1 | 0.2 |
| 7 | Group C&D Adrenoleukodystrophy CBL Pre Test (out of 10) | 2.7 | 1.6 | 0.2 |
| 8 | Group C&D Adrenoleukodystrophy CBL Post Test (out of 10) | 7.4 | 1.2 | 0.2 |
| 9 | Group A&B Formative Assessment Pompe (out of 5) | 2.0 | 1.3 | 0.2 |
| 10 | Group C&D Formative assessment Pompe (out of 5) | 3.0 | 1.3 | 0.2 |
| 11 | Group A&B Formative Assessment Other (out of 95) | 41.9 | 11.9 | 1.6 |
| 12 | Group C&D Formative assessment Other (out of 95) | 51.8 | 12.7 | 2.1 |
| 13 | Group A&B Formative Assessment Before Intervention (out of 100) | 55.4 | 11.7 | 1.7 |
| 14 | Group C&D Formative assessment Before Intervention (out of 100) | 57.9 | 10.1 | 1.8 |

Supplementary Table 3

Themes identified through content analysis of learner’s feedback and facilitators FGD

| Theme | Agreement | Partial agreement | Disagreement |
| --- | --- | --- | --- |
| Cinemeducation was fun, interesting, and engaging | Using movie clips make case-based learning more interesting and fun. | “I couldn't see any drawbacks. But preventing sleeping tendency, it's better to provide coffee.”  *“I felt excited to see the enthusiasm and interest of the students during the teaching-learning process. However, the student’s perceptions of the movies were somewhat different than what was expected. Though Lorenzo`s Oil had more components of Biochemistry, students liked Extraordinary Measures more because of the star performers like Harrison Ford.* |  |
| Cinemeducation was empathizing, relatable, and lifelike | “The medical cases become more relatable with real-life scenarios.”  “As we empathized with the patient, we became more enthusiastic to learn about the pathogenesis and management approaches.”  “Apart from learning the method, it is much more important how to behave with the patients suffering from the disease. As future doctors, we should give hope and try our best to save a patient’s life.” | “Though only relevant clips and not the full movie was shown still we had a positive emotional influence depending on the skills of the actors.” | “There was no emotional impact”. |
| Cinemeducation led to better correlation, visualization, and insight | “Movies provided background information regarding the natural history of the disease which helped us in visualizing the Biochemical concepts and mechanistic insights.”  “We were able to better correlate Biochemical knowledge with clinical situations.” |  |  |
| Cinemeducation increased attention span, interactivity, and thinking | “This is an innovative way of arousing interest in us resulting in increased attention span and increased retention of the concepts.”  “This is an innovative approach to provoke thinking in the students leading to accelerated learning and better retention.” |  |  |
| Cinemeducation increased the satisfaction of learners and facilitators | “Movie clips would be a more effective learning tool than conventional CBL when it is used to supplement the interactive mode of teaching.” |  |  |
| There was a dearth of suitable movies | *“The biggest hurdle would be the dearth of suitable movies for teaching Biochemistry.”*  *“As the limited number of movies are available for teaching Biochemistry, we need to make small movies ourselves. We may rope in professionals and also interested students may be from Second or Third Professional Years who have received some clinical exposure.”* |  |  |
| Foreign language and accent were problematic | “According to me, one drawback Is there, that in movie clips there are mostly Hollywood movies (That means English speaking). But problem is that I can't understand that hard language so at best try to put subtitle of the language below.”  “The unfamiliar accents in some movies would require subtitles to be added.”  *“The language is a barrier for students from vernacular mediums also the Italian accent in Lorenzo`s Oil created problems for the learners. The English subtitle was needed to be added in all the clips.”* |  |  |
| Cinemeducation was time-consuming | “Moreover, using movie clips can consume a lot of time it might be challenging to retain the depth of coverage of a topic.” |  |  |
| Cinemeducation led to less coverage of the subject | “As all aspects of a topic would usually not be covered in a movie, some information might be missing. Some students would be indulging more in the characters rather than the topic. After showing movies a lot of discussions would be required to make the learning effective.” |  |  |
| Movies had incorrect or missing information | *“This is also important to add narration while showing the movie clips since some of the information provided in the movie may not be scientifically correct like in the last scene of Extraordinary Measures where the children receiving enzyme replacement therapy were giggling which was attributed to Glycogen breaking down to Glucose in muscle.”* |  |  |

Supplementary figure 1

Q - Q plot

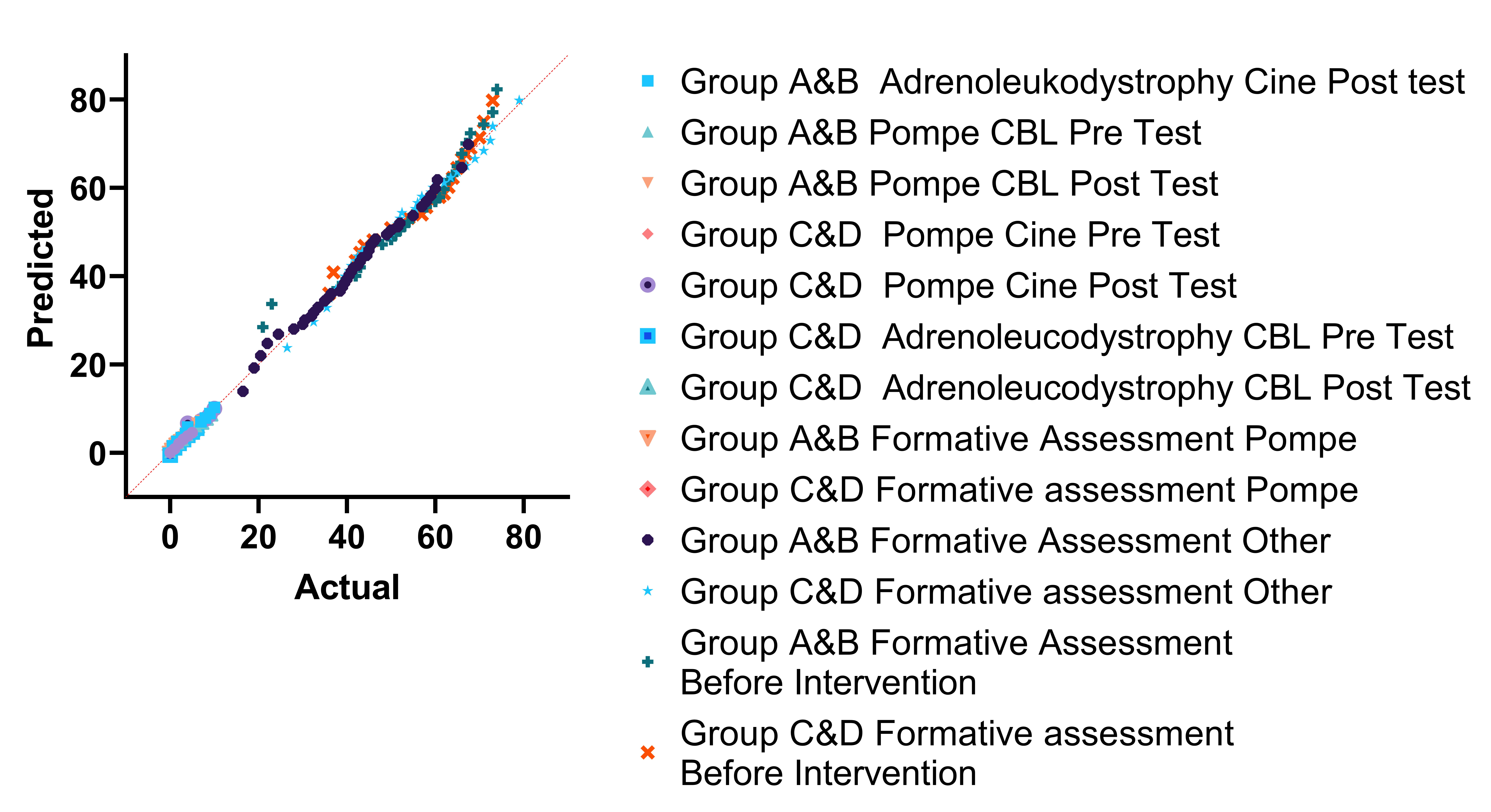
*Supplementary figure 1: Both the pre-test, post-test, and formative assessment scores for the four groups were found to be not normally distributed. Hence nonparametric tests were used for further data analysis.*
